# AAV-mediated progranulin delivery to a mouse model of progranulin deficiency causes T cell-mediated hippocampal degeneration

**DOI:** 10.1101/308692

**Authors:** Defne A. Amado, Julianne M. Rieders, Fortunay Diatta, Pilar Hernandez-Con, Adina Singer, Junxian Zhang, Eric Lancaster, Beverly L. Davidson, Alice S. Chen-Plotkin

**Affiliations:** Department of Neurology, Perelman School of Medicine, University of Pennsylvania, Philadelphia, PA USA 19104; Department of Pathology and Laboratory Medicine, University of Pennsylvania, Philadelphia, PA USA 19104; Children’s Hospital of Philadelphia, Philadelphia, PA USA 19104

## Abstract

Adeno-associated virus (AAV)-mediated gene replacement is emerging as a safe and effective means of correcting single-gene mutations, and use of AAV vectors for treatment of diseases of the CNS is increasing. AAV-mediated progranulin gene (*GRN*) delivery has been proposed as a treatment for *GRN*-deficient frontotemporal dementia (FTD) and neuronal ceroid lipofuscinosis (NCL), and two recent studies using focal intraparenchymal AAV-*Grn* delivery to brain have shown moderate success in histopathologic and behavioral rescue in mouse FTD models. Here, we used AAV9 to deliver *GRN* to the lateral ventricle to achieve widespread expression in the *Grn* null mouse brain. We found that despite a global increase in progranulin throughout many brain regions, overexpression of *GRN* resulted in dramatic and selective hippocampal toxicity and degeneration affecting both neurons and glia. Histologically, hippocampal degeneration was preceded by T cell infiltration and perivascular cuffing, suggesting an inflammatory component to the ensuing neuronal loss. *GRN* delivery with an ependymal-targeting AAV for selective secretion of progranulin into the cerebrospinal fluid (CSF) similarly resulted in T cell infiltration as well as ependymal hypertrophy. Interestingly, overexpression of *GRN* in wild-type animals also provoked T cell infiltration. These results call into question the safety of *GRN* overexpression in the CNS, with evidence for both a region-selective immune response and cellular proliferative response following *GRN* gene delivery. Our results highlight the importance of careful consideration of target gene biology and cellular response to overexpression in relevant animal models prior to progressing to the clinic.

**Significance Statement:** Gene therapies using adeno-associated viral (AAV) vectors show great promise for many human diseases, including diseases that affect the central nervous system (CNS). Frontotemporal dementia (FTD) and neuronal ceroid lipofuscinosis (NCL) are neurodegenerative diseases resulting from loss of one or both copies of the gene encoding progranulin (*GRN*), and gene replacement has been proposed for these currently untreatable disorders. Here, we used two different AAV vectors to induce widespread brain *GRN* expression in mice lacking the gene, as well as in wild-type mice. Unexpectedly, GRN overexpression resulted in T cell infiltration, followed by marked hippocampal neurodegeneration. Our results call into question the safety of GRN overexpression in the CNS, with wider implications for development of CNS gene therapies.

## Introduction

Frontotemporal dementia (FTD) (1, 2) and neuronal ceroid lipofuscinosis (NCL) (3) are neurodegenerative diseases resulting from haploinsufficiency or complete deficiency of progranulin (GRN), which is encoded by the gene *GRN*. FTD manifests in late middle age, with symptoms ranging from behavioral changes to deterioration of language and with death ensuing in a mean of 3-5 years after diagnosis (4). Mutations in *GRN* are a highly penetrant cause of FTD and account for up to 25% of inherited FTD cases (1, 2). Nearly 70 *GRN* mutations have been identified that cause FTD, and >90% are nonsense mutations that lead to *GRN* haploinsufficiency (1, 2, 5), with the remainder resulting in functional haploinsufficiency through other means (*e.g*. defects in GRN secretion). For reasons that are still poorly understood, *GRN*-deficient states result in accumulation of Tar-DNA binding protein of 43kD (TDP-43) (1, 2), in characteristic inclusion bodies, with subsequent neuronal loss and atrophy of the frontal and temporal lobes.

In the case of NCL, complete *GRN* deficiency leads to lysosomal dysfunction and accumulation of lipofuscin in neurons and other cell types and a clinical syndrome consisting of generalized seizures, mild cognitive dysfunction, vision loss, cerebellar degeneration, and palinopsia (6–8). Strategies to boost GRN in the FTD and NCL brain have been under development since its discovery as a major causal mutation for these diseases (9-11).

GRN is a secreted growth factor involved in embryonic development, wound healing, and immune modulation (12, 13). In the mouse brain, *Grn* is expressed highly in mature neurons and microglia and is upregulated in activated microglia following injury (14). In human postmortem brain tissue, GRN expression is widespread in both normal and FTD subjects (15). Both *in vitro* and *in vivo*, GRN has been shown to play a role in neuronal survival and neurite outgrowth (16-18), and a neuronal receptor for GRN, sortilin, has been identified (19). Based on its growth promoting properties, augmentation of GRN has been considered for the treatment of a wide range of neurodegenerative diseases. Indeed, lentivirus- and AAV-mediated *GRN* delivery to the central nervous system (CNS) have been investigated in preclinical models of Alzheimer’s disease (20, 21), Parkinson’s disease (22), motor neuron disease (23, 24), and Huntington’s disease (25).

Methods to augment or replace *GRN* expression include activating transcription (10), enhancing translation (9), increasing the levels of extracellular GRN (11), or using gene therapy based approaches. Among the latter, gene delivery using adeno-associated viral (AAV) vectors has risen to the forefront of gene therapy on the basis of its excellent safety and efficacy profile. AAV-mediated gene delivery has been used successfully in preclinical models for several decades and has recently shown success in treating a diverse range of diseases in humans, including hemophilia, Leber’s Congenital Amaurosis, and spinal muscular atrophy (26-28). To achieve CNS transduction, AAV can be delivered to the brain parenchyma or the cerebrospinal fluid (CSF), either via intrathecal delivery or by injection into the lateral ventricle, with therapeutic benefit in preclinical models with both gain-of-function and loss-of-function diseases (29-34). Interestingly, and in contrast to peripherally administered AAV (35, 36), numerous studies using different AAV vectors with various gene targets have shown minimal innate or adaptive immune response to AAV-mediated gene delivery in the CNS, even in animals with no prior exposure to the gene in question.

A recent study using direct bilateral injection of AAV1.*Grn* into the medial prefrontal cortex (mPFC) of *Grn* null mice demonstrated improvements in lipofuscinosis and microgliosis, with focal improvements in lysosomal function (37). This group had previously used this approach in *Grn* haploinsufficient mice and demonstrated improvement in lysosomal readouts and social dominance deficits (38). Notably, AAV *GRN* gene therapy in *Grn* null mice displayed robust microglial activation at the injection site, with induction of anti-GRN antibodies (37). No other immunologic phenotypes were reported in this short-term study.

While these studies are promising, translation of intraparenchymal gene delivery to the human brain is a challenge based on the size of the target compared to the murine brain. The aim of our study was to deliver *GRN* globally using a method easily translatable to human subjects, namely a single intraventricular injection of AAV.*GRN*. As a first approach, we selected AAV9 on the basis of its ability to broadly disperse and infect neurons and glia after CSF delivery, as well as its track record of use in prior studies (28, 34, 39). We also tested AAV4 due to its selectivity for ependymal cells and excellent safety profile (30, 40, 41), to maximize CSF secretion with the goal of broad CNS uptake through the neuronal receptor sortilin. Regardless of serotype, our studies show that over-expression of *GRN* in brain for extended periods of time is deleterious, causing profound neurodegeneration and raising concern about excessive expression of *GRN* in mammalian brain as a therapy for FTD/NCL.

## Results

### Characterization of the *Grn* Null Phenotype

Mice lacking *Grn* have an age-dependent histopathologic phenotype consisting of habenular and hippocampal vacuolation and increased ubiquitination starting at 7 months of age, as well as diffusely increased astrogliosis and microgliosis starting at 12 months of age (42-44). In our *Grn* null animals, we confirmed the previously reported increase in vacuolation (42), which was most pronounced in the habenula and increased with age (**Fig. 1*A***, arrowheads). Additionally, we noted astrocytosis in the *Grn* null striatum that is present as early as 6 months and progresses with age; this histopathological finding was not present even in 12-month-old WT mice (**Fig. 1*B***) and has not been previously described. Hippocampal morphology was unaffected by genotype at any age (**Fig. 1*C***).

**Fig. 1.**
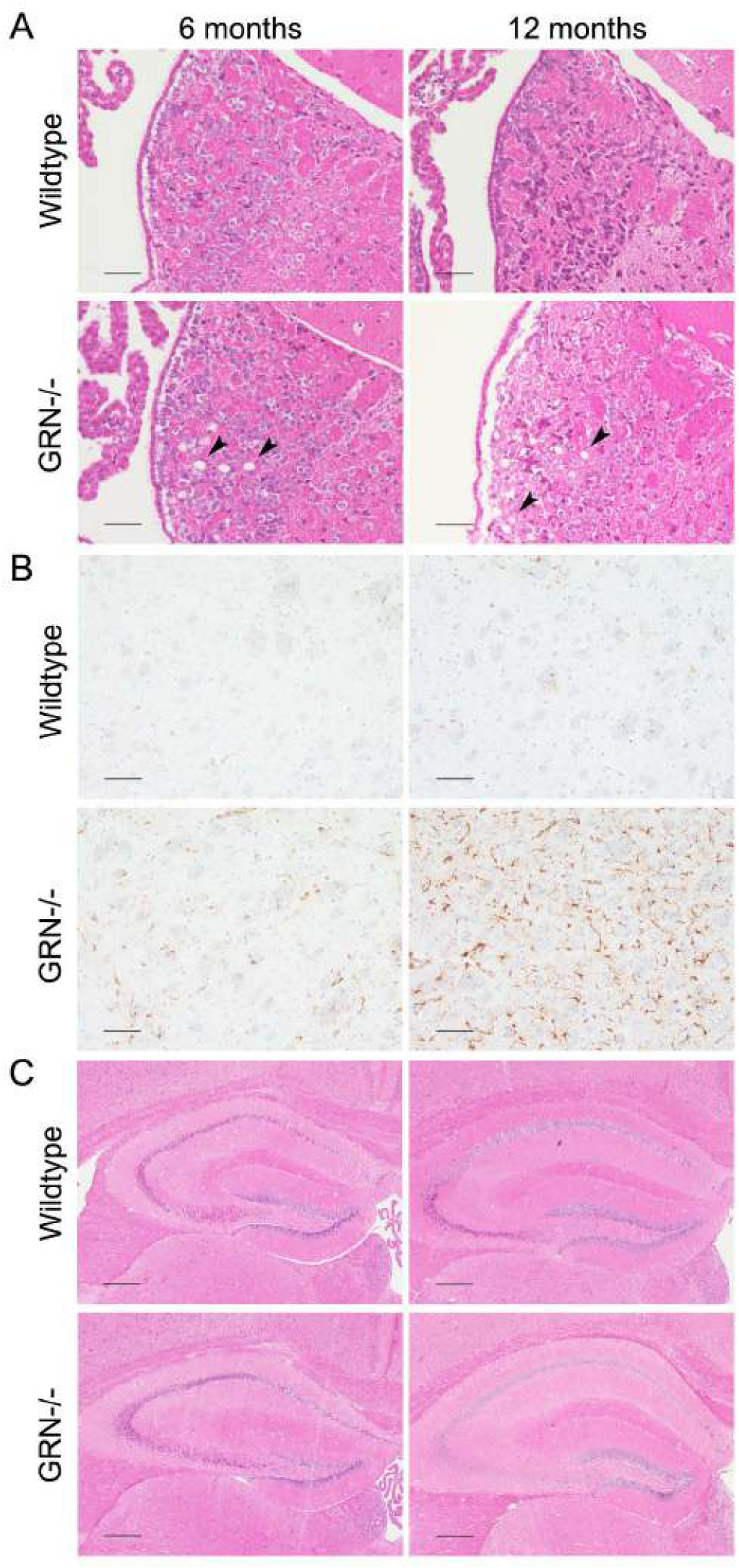
*Grn* null mice recapitulate previously published histopathologic findings and exhibit previously undescribed abnormalities. (A) *Grn* null mice exhibit vacuolation that is most pronounced in the habenula and increases with age (arrowheads), and is absent from wildtype mice at all time points. (Scale bar: 50 μm.) (B) *Grn* null mice demonstrate an age-dependent increase in astrocytosis compared to wildtype mice, as seen by GFAP staining. Shown here is the striatum, an area in which astrocytosis in *Grn* null mice has not been previously described. (Scale bar: 100 μm.) (C) The hippocampus shows no gross morphological differences in *Grn* null mice compared to WT at 6 or 12 months. (Scale bar: 250 μm.)

### AAV-Mediated Gene Transfer Results in Sustained GRN Expression in *Grn* Null Mice

*Grn* null mice were injected with AAV9 encoding human *GRN* (AAV9.*hGRN*) (Fig. S1) into the right posterior lateral ventricle at 6-7.5 months of age and sacrificed at time points ranging from 1 to 6 months post-injection as indicated (**Fig. 2*A***). We observed the highest levels of GRN expression, as detected by enzyme-linked immunoassay (ELISA), in the ipsilateral periventricular region; GRN levels remained undiminished at 1-, 3-, 4.5- and 6 months post-injection compared to uninjected whole brain tissue from null mice (**Fig. 2*A***, left). We also observed high GRN levels in the ipsilateral striatum and to a lesser extent, the ipsilateral frontal cortex, although cortical expression diminished over time. Similarly high levels of GRN were detected in the contralateral periventricular region (**Fig. 2*A***, right), but with minimal increase in the left striatum or left frontal cortex. GRN was additionally detected at moderate levels in the periventricular region of the 3rd ventricle, the brainstem, and the spinal cord at all time points (Fig. S2). These data indicate broad, sustained *hGRN* expression.

**Fig. 2.**
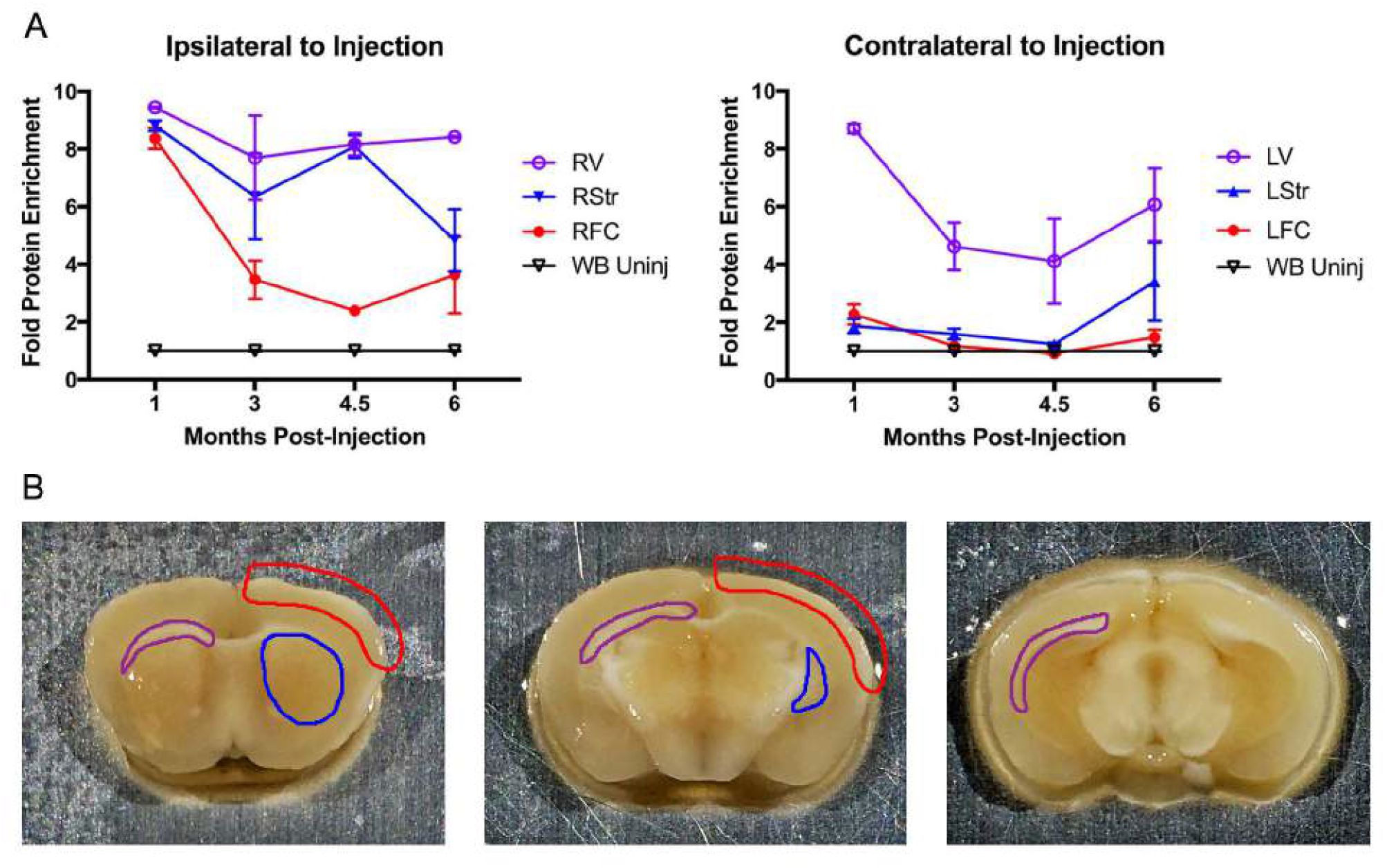
AAV9 mediates sustained expression of GRN in *Grn* null mouse brain. *Grn* null mice were injected at 6-7.5 months of age with AAV9.*hGRN* or AAV9.*eGFP* in the right lateral ventricle and sacrificed 1, 3, 4.5, or 6 months post-injection. Brains were microdissected and GRN levels measured by ELISA. (A, left) GRN levels in the right (injected) peri-ventricular area (RV), right striatum (RStr), right frontal cortex (RFC) and uninjected homogenized whole brain (WB) are shown. (A, right) GRN levels in the left (uninjected) peri-ventricular area (LV), left striatum (LStr), left frontal cortex (LFC) and uninjected homogenized whole brain (WB) are shown. (B) Schematic illustrating the regions collected by microdissection in blue (striatum), red (cortex), and purple (peri-ventricular area). *n* = 3 mice/group at each time point.

### Overexpression of *GRN* Results in Progressive Hippocampal

Having established sustained expression of *GRN*, we next performed detailed histological and immunohistochemical analyses to assess for rescue in our treated mice. In *Grn* null animals injected with AAV9.*hGRN* at 6-7.5 months of age and sacrificed 6 months after injection, we found striking morphological changes in the hemisphere ipsilateral to injection. Specifically, in the majority of injected animals, the hippocampus ipsilateral to the injection site demonstrated marked loss of structural integrity, while adjacent structures as well as the contralateral hippocampus appeared relatively unaffected (**Fig. 3*A***). Across all affected mice, the most prominent and consistent changes involved the inferomedial hippocampus, and especially the dentate region (**Fig. 3*A***, bottom, and **Fig. 3*B***, right panels). Similar hippocampal degeneration occurred in *Grn* null mice injected with an AAV9 vector delivering mouse *Grn* (Fig. S3), indicating that the response was not specific to delivery of a human gene.

**Fig. 3.**
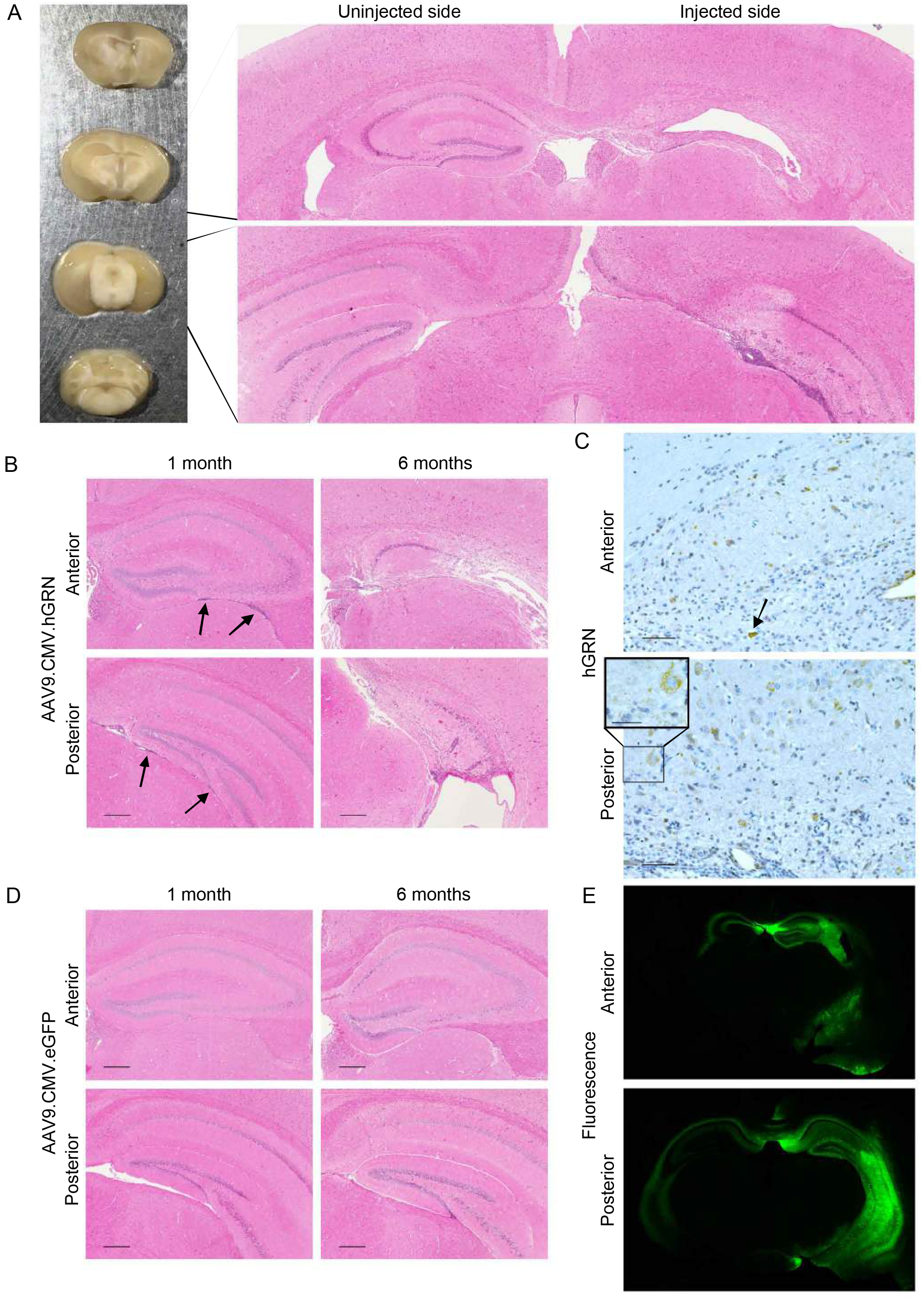
Overexpression of GRN is toxic to cells of the hippocampus. Mice were injected at 6-7.5 months of age and sacrificed 1-6 months post-injection, and brains were either embedded and stained for immunohistochemical analysis (A-D) or sectioned and imaged fluoroscopically (E). (A, left) Gross morphological differences were observed between the injected and uninjected hemispheres in 6 of 11 mice 6 months after injection of AAV9.*hGRN*, with marked atrophy of the hippocampus on the injected side. (A, right) H&E of AAV9.*hGRN* injected mouse brain reveals marked degeneration of the hippocampus ipsilateral to injection and a dense cellular infiltrate throughout the remaining hippocampal tissue, extending from anterior (upper panel) to posterior (lower panel). (B) The cellular infiltrate is observed along the length of the hippocampus as early as 1 month post-injection of AAV9.*hGRN*, both inferior to (arrows) and within the parenchyma of the hippocampus (left panels; *n* = 3 mice/group). By 6 months, marked morphological changes are observed (right panels). (Scale bars: 250 μm.) (C) High levels of GRN are detected throughout the hippocampus at all time points, with a diffuse cytoplasmic pattern of expression (inset). In some cases, cells expressing GRN exhibit pyknotic nuclei (arrow). Shown here is expression 6 months post injection. (Scale bars: 50 μm; inset: 25 μm.) (D) AAV9.*eGFP* injected brains appear normal at 1- and 6 months post injection (*n* = 1 and 4 mice/group respectively) and are similar in appearance to those of uninjected *Grn* null mice (*n* = 4, data not shown). (Scale bars: 250 μm.) (E) High levels of *eGFP* expression are detected by fluorescence microscopy throughout the hemisphere ipsilateral to injection, with some cross-over to the contralateral side. Shown here is expression 3 months post-injection (*n* = 2 mice).

To better define the timeline of degenerative changes, animals were harvested soon after injection. At one month post-injection with AAV9.*hGRN*, a hypercellular infiltrate was noted to extend anteriorly and posteriorly along the entire hippocampus (**Fig. 3*B***, left panels), most prominent inferior to and within the hippocampal parenchyma. By 6 months post-injection, prominent hypercellular infiltrates and perivascular cuffing accompanied loss of recognizable hippocampal structures (**Fig. 3*B***, right panels). Staining confirmed strong GRN expression in these regions at all time points (**Fig. 3*C***). Positive staining was noted in neurons that appeared to be healthy (**Fig. 3*C***, lower inset) and in cells with pyknotic nuclei (**Fig. 3*C***, arrow). In contrast, littermate control animals treated with AAV9-delivered *eGFP* (AAV9.*eGFP*) showed no pathology at 1- or 6 months post injection (**Fig. 3*D***) and appeared similar to uninjected littermates (data not shown), despite high levels of eGFP expression (**Fig. 3*E***).

### Responses to *hGRN* Overexpression are Region and Cell-Type Specific

Gross morphological assessments were performed to determine if the degeneration observed after AAV9.*hGRN* delivery was present in other brain regions with high levels of expression. No gross morphological differences were seen between the cortex ipsilateral to the injection site and either the contralateral cortex or that of AAV9.*eGFP*-injected controls at 6 months post-injection (**Fig. 4*A-B***), despite moderately high GRN levels in cortical brain isolates (**Fig. 2**). Indeed, cortical neuron organization remained intact ipsilateral to the site of AAV9.*hGRN* injection (**Fig. 4*C***), and no obvious changes in architecture, neuronal number, or gliosis were found in the ipsilateral striatum (**Fig. 4*D-G***), an area with high GRN levels (**Fig. 2**). Brain tissue from AAV9.*eGFP*-injected and uninjected animals was similar in appearance across all parameters (data not shown). These results suggest that *hGRN* overexpression selectively affects hippocampal brain regions.

**Fig. 4.**
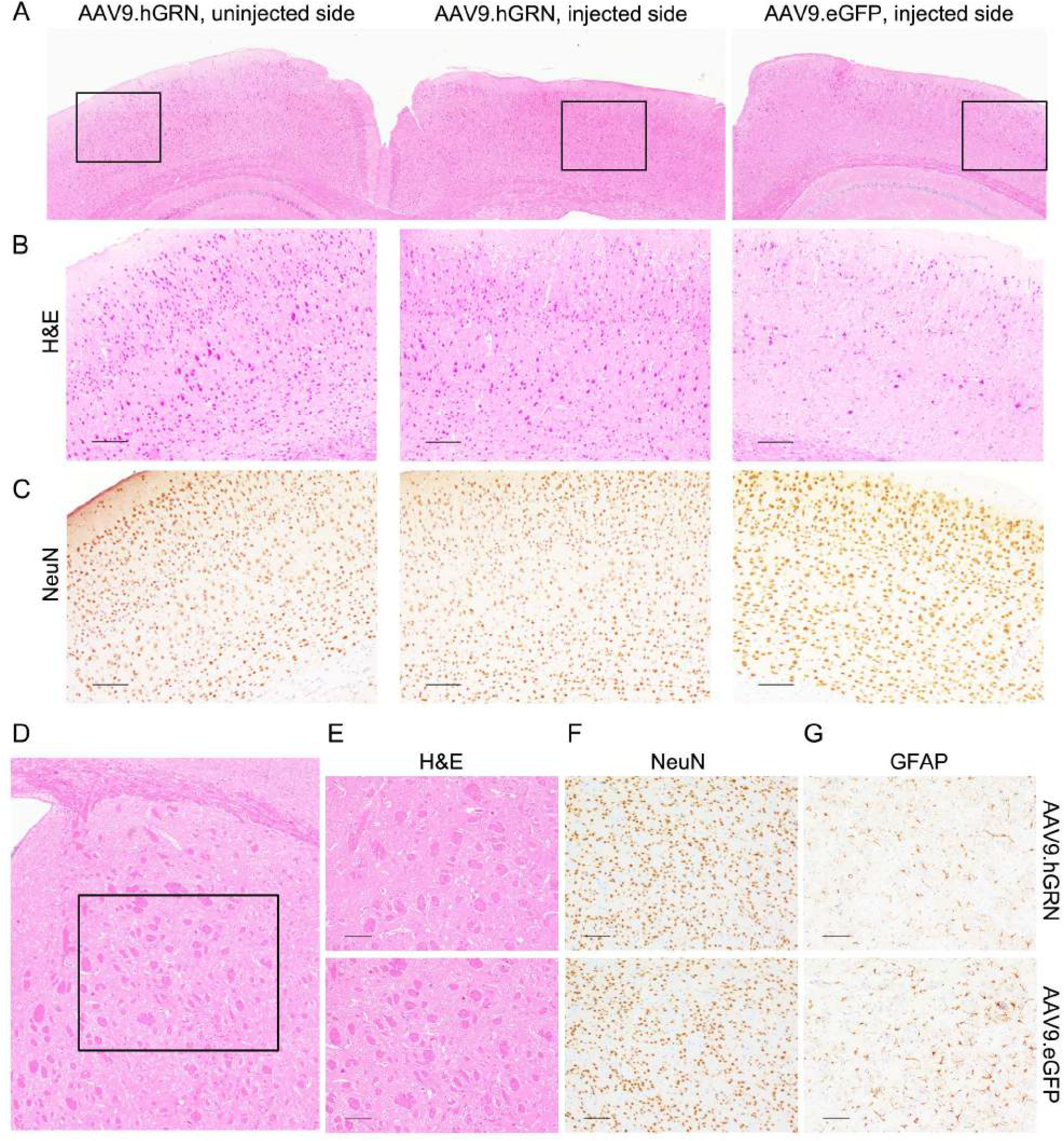
The cortex and striatum are unaffected by GRN overexpression. (A) The ipsilateral cortex immediately adjacent to the degenerated hippocampus appears unaffected by AAV9.*hGRN* overexpression when compared to the contralateral cortex and to cortex ipsilateral to AAV9.*eGFP* injected brain. Gross morphology, layer organization, and neuronal numbers appear unremarkable by H&E (B) and NeuN staining (C). (Scale bars: 100 μm.) Similarly, the ipsilateral striatum directly anterior to the degenerated hippocampus (D) appears unremarkable by H&E (E), NeuN (F), and GFAP (G) staining compared to AAV9.*eGFP* injected controls 6 months post-injection (scale bars: 100 μm), as well as to uninjected controls (data not shown). *n* = 11 AAV9.hGRN-injected mice/group, *n* = 4 AAV9.*eGFP*-injected mice/group.

To determine what cell types were affected in the hippocampus anteriorly (**Fig. 5*A-D***) and posteriorly (**Fig. *5E-H***), tissue sections were stained for the neuronal marker NeuN, the glial marker GFAP, and the microglial marker Iba-1. Across all affected mice, extensive hippocampal neuronal loss was noted, with the dentate gyrus most severely affected (**Fig. 5*B*** and **5*F***). Hippocampal astrocytes demonstrated fewer, less robust processes (**Fig. 5*C*** and **5*G***), and microglial infiltration ipsilateral to injection was prominent, particularly in the posterior hippocampus (**Fig. 5*D*** and **5*H***) (45). In all cases, the side contralateral to the injection (left panels) appeared similar to the hippocampus of both AAV9.*eGFP*-injected and uninjected control littermates (data not shown). These data indicate that AAV9-mediated *hGRN* overexpression is toxic to neurons and astrocytes in the hippocampus, and provokes a strong local microglial response.

**Fig. 5.**
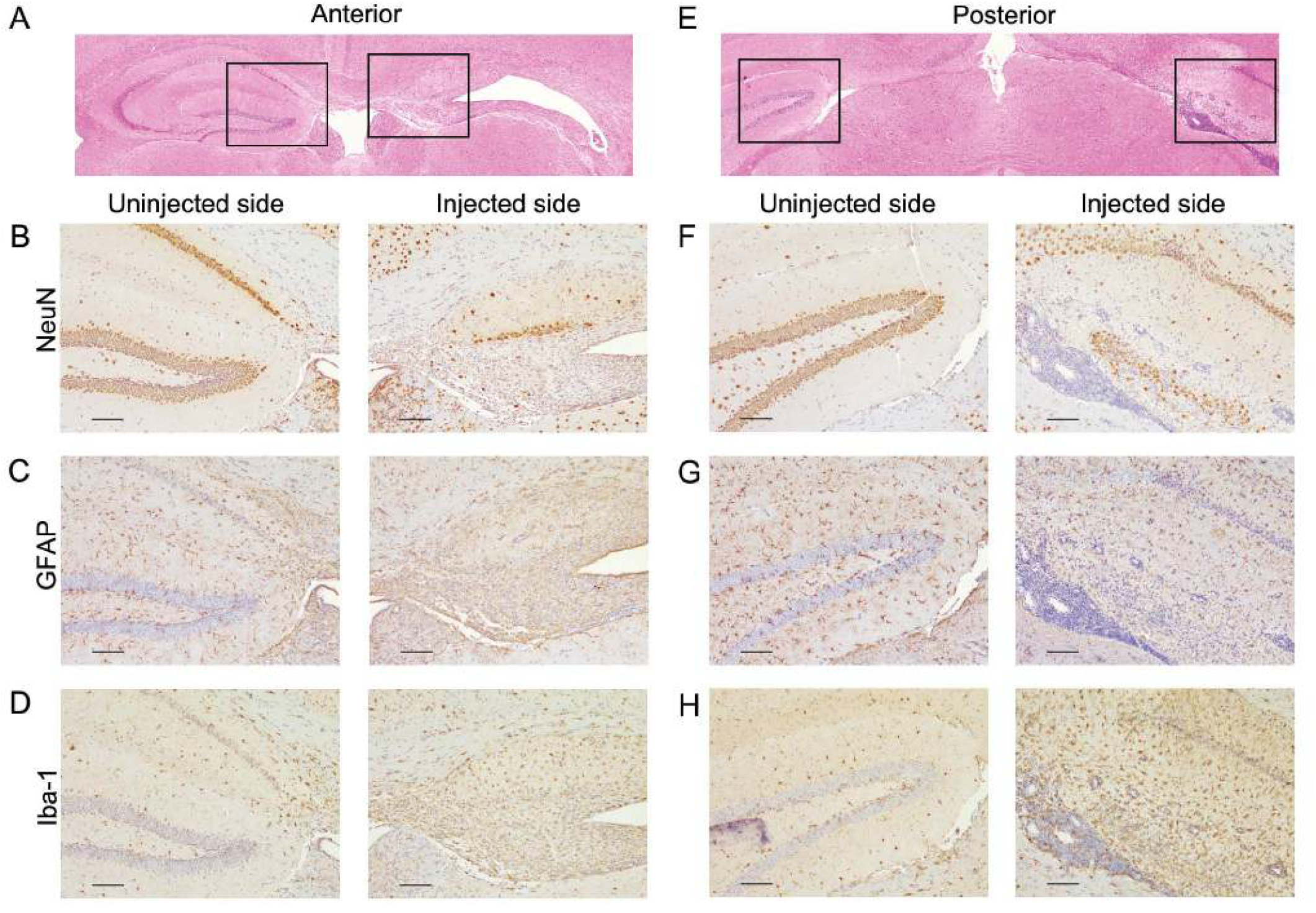
AAV9.*hGRN*-overexpressing mice undergo cell-specific hippocampal degeneration. H&E-stained coronal sections show hippocampal degeneration 6 months post-injection on the injected side anteriorly (A) and posteriorly (E), with boxes indicating regions magnified below (B-D, F-H). On the injected side, NeuN staining shows striking neuronal loss throughout all regions of the hippocampus both anteriorly (B) and posteriorly (F); GFAP staining shows loss of astrocytic processes in anterior (C) and posterior (G) hippocampus; Iba-1 staining for microglia shows a dense microglial infiltrate in the hippocampal region that is present anteriorly (D) but is most prominent posteriorly (H). Throughout the posterior hippocampus there is a dense cellular infiltrate in the ependymal space underlying the hippocampus (F-H) ipsilateral to injection. (Scale bars: 100 μm.)

### A T Cell Mediated Inflammatory Response Precedes Neuronal Loss and Occurs in Both Wild-Type and *Grn* Null Animals

The hypercellular infiltrate found in AAV9.*hGRN*-injected animals consists of cells with a high nucleus-to-cytoplasm ratio characteristic of lymphocytes. As such, sections were stained for the cell proliferation marker Ki-67, the B lymphocyte marker B220 (CD45), and the T lymphocyte marker CD3.

As shown in **Fig. 6**, abundant proliferative cells were noted (**Fig. 6*B*** and **6*F***), and the majority of these cells were positive for CD3 (**Fig. 6*C*** and **6*G***). In both anterior and posterior sections, there was extensive perivascular cuffing by CD3+ cells both within and adjacent to the hippocampus (**Fig. 6*G***, arrowheads). These data collectively indicate a robust T cell infiltration of the hippocampal region with minimal contribution from B cells.

**Fig. 6.**
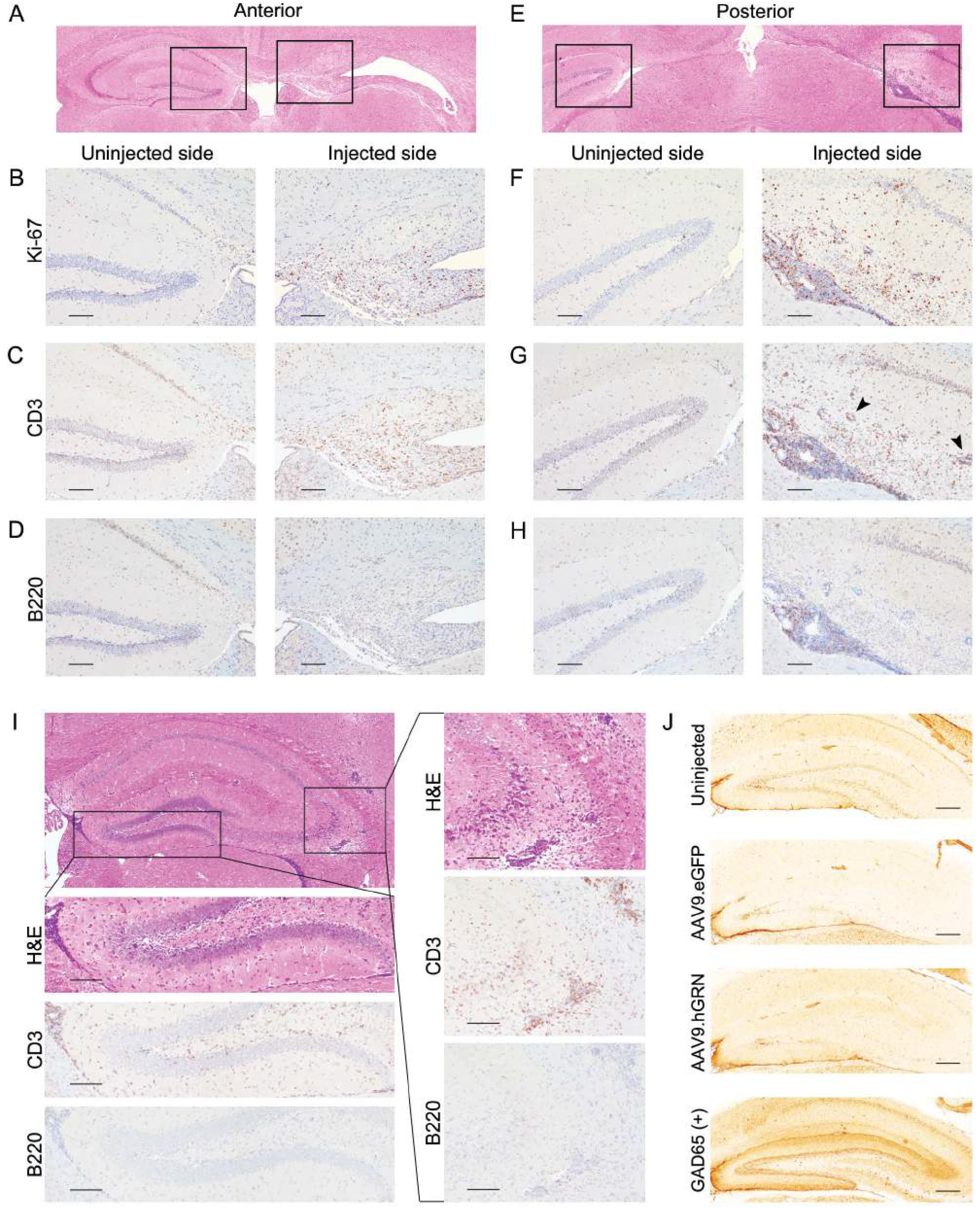
Hippocampal degeneration is characterized by a T cell inflammatory response that precedes cell death. Mice were injected with AAV9.*hGRN* at 6-8 months of age and sacrificed 6 months post-injection (A-H) or 1 month post-injection (I). H&E-stained coronal sections show hippocampal degeneration 6 months post-injection on the injected side anteriorly (A) and posteriorly (E), with boxes indicating regions magnified below (B-D, F-H). On the injected side, Ki-67 staining demonstrates proliferating cells throughout the hippocampus anteriorly (B) and more markedly posteriorly (F); CD3 staining identifies most of these proliferating cells as T cells (C, G), while B220 staining for B cells is largely negative (D, H), aside from some positively stained cells in the dense posterior sub-hippocampal infiltrate (H). (Scale bars: 100 μm.) Mice were then examined at 1 month post injection (I). H&E staining shows a dense infiltrate in the ependymal space inferior to the hippocampus that is CD3+ and B220− (left box and magnifications below; scale bars: 100 μm). In the CA2/3 region, there is also dense hypercellularity with perivascular cuffing that is CD3+ and B220- (right box and magnifications on right; scale bars: 100 μm, *n* = 3 mice). (J) Rat brain slices were incubated with mouse serum from uninjected (top, *n* = 3), AAV9.*eGFP*-injected (second, *n* = 2), or AAV9.*hGRN*-injected (third, *n* = 5) mice at 6 months post injection, with GAD65+ mouse serum used as a positive control for antibody-based hippocampal reactivity (bottom). There was no immune reactivity in serum collected from mice injected with AAV9.*hGRN* at 6 months, or at 1- (*n* = 3) or 3 (*n* = 2) months after AAV9.*hGRN* injection (data not shown). (Scale bars: 250 μm.)

To test if inflammatory infiltrates precede or follow the hippocampal degeneration, brain sections from animals sacrificed at earlier time points after AAV9.*hGRN* injection were characterized. At one month post-injection, hippocampal structures were maintained despite dense cellular infiltration ventral to the hippocampus, with widespread perivascular cuffing lateral to and within the hippocampus (**Fig. 6*I***). Infiltrating cells were positive for CD3 and negative for B220, indicating that T cell infiltration precedes hippocampal neurodegeneration.

In human autoimmune encephalitides, hippocampal degeneration often ensues from autoantibodies specific for hippocampal antigens. Triggering events may be expression of an ectopic antigenic protein by a tumor, or unmasking of a native hippocampal antigen by an inflammatory process (46). We thus tested whether *hGRN* overexpression elicited a similar pathophysiological process in our *Grn* null animals using a previously-described rat hippocampal slice assay (47). As shown in **Fig. 6*J***, serum from mice with the hippocampal degeneration phenotype screened negative for anti-hippocampal antibodies, regardless of whether serum was drawn 1-, 3-, or 6 months after AAV9.*hGRN* injection. Moreover, AAV9.*hGRN* intraventricular delivery into wild-type animals also elicited perivascular cuffing with infiltration of CD3 positive T cells as early as one month after gene delivery, which became prominent and was accompanied by loss of hippocampal structures by 3 months post-injection (**Fig. 7**). Taken together, these data support a role for T cell mediated hippocampal neurodegeneration following *GRN* overexpression.

**Fig. 7.**
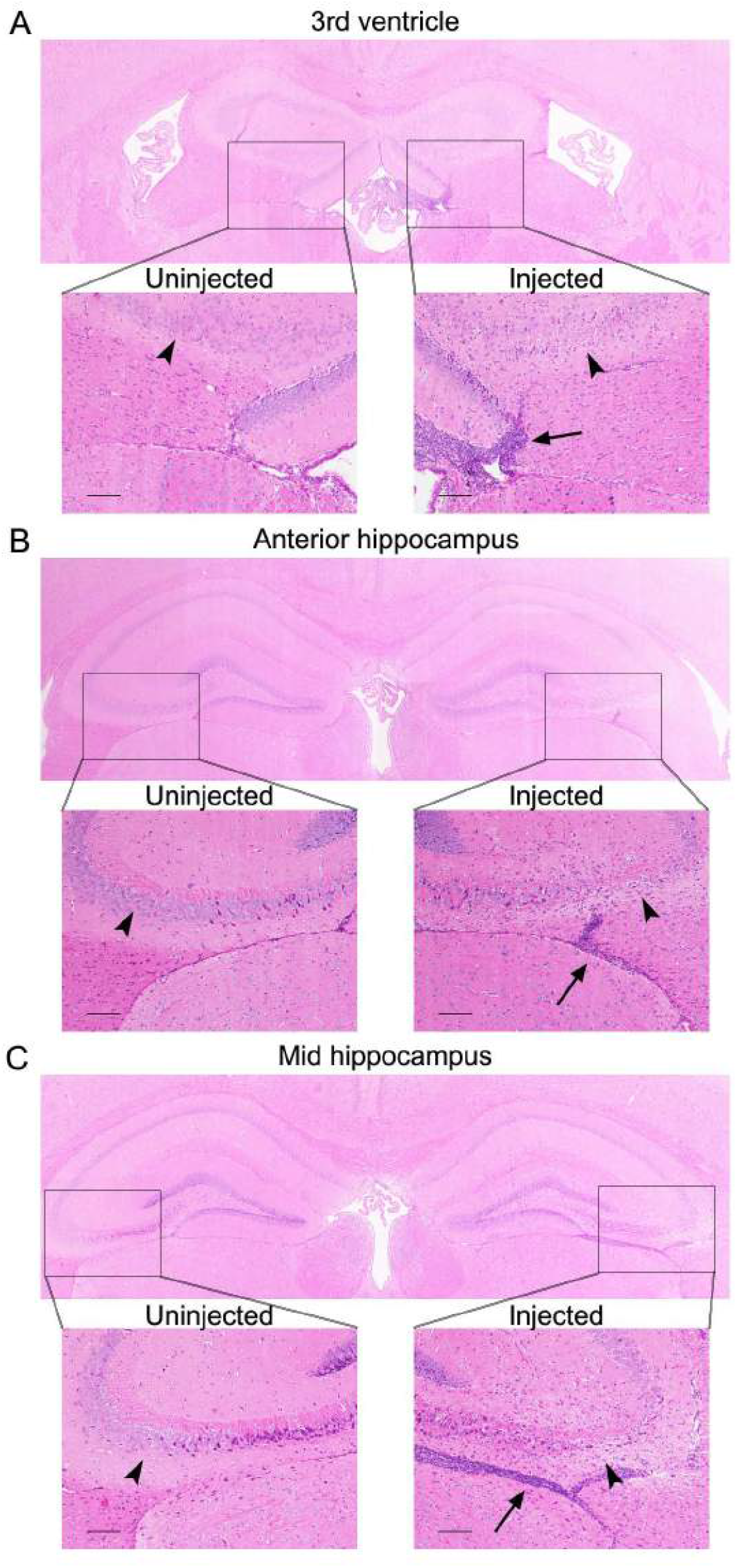
Wildtype mice also mount a T cell response accompanied by hippocampal cellular loss after injection with AAV9.*hGRN*. Wildtype background-matched mice were injected in the right lateral ventricle at 6 months of age with AAV9.*hGRN* and sacrificed 1- (data not shown) or 3 months post-injection. A hypercellular infiltrate was observed most prominently adjacent to the third ventricle ipsilateral to injection (A) and extending posteriorly throughout the hippocampus (B, C). The infiltrate extended inferior to the hippocampus (arrows) as well as within the parenchyma, where we observed marked loss of cells in the CA2/3 region of the hippocampus on the injected side compared to the uninjected side (arrowheads; *n* = 3 mice).

### *hGRN* Delivered by the AAV4 Ependymal-targeting Vector Elicits an Inflammatory Response and Ependymal Hypertrophy

AAV9 transduces multiple cell types, including neurons and glia (48). We next asked whether the observed inflammatory response and subsequent hippocampal degeneration was serotype specific. For this we used AAV4, an ependymal-targeting serotype (49) that allows secretion of transgene products into the CSF (30, 41, 50). In *Grn* null mice injected with AAV4.*hGRN* 1 month earlier into the right lateral ventricle, at the same dose and age as our AAV9.*hGRN*-injected animals, we observed a hypercellular infiltrate in the anterior (**Fig. 8*A-B***) and posterior (**Fig. 8*C***) hippocampus. Consistent with our previous findings, infiltrating cells were positive for the T cell marker CD3.

**Fig. 8.**
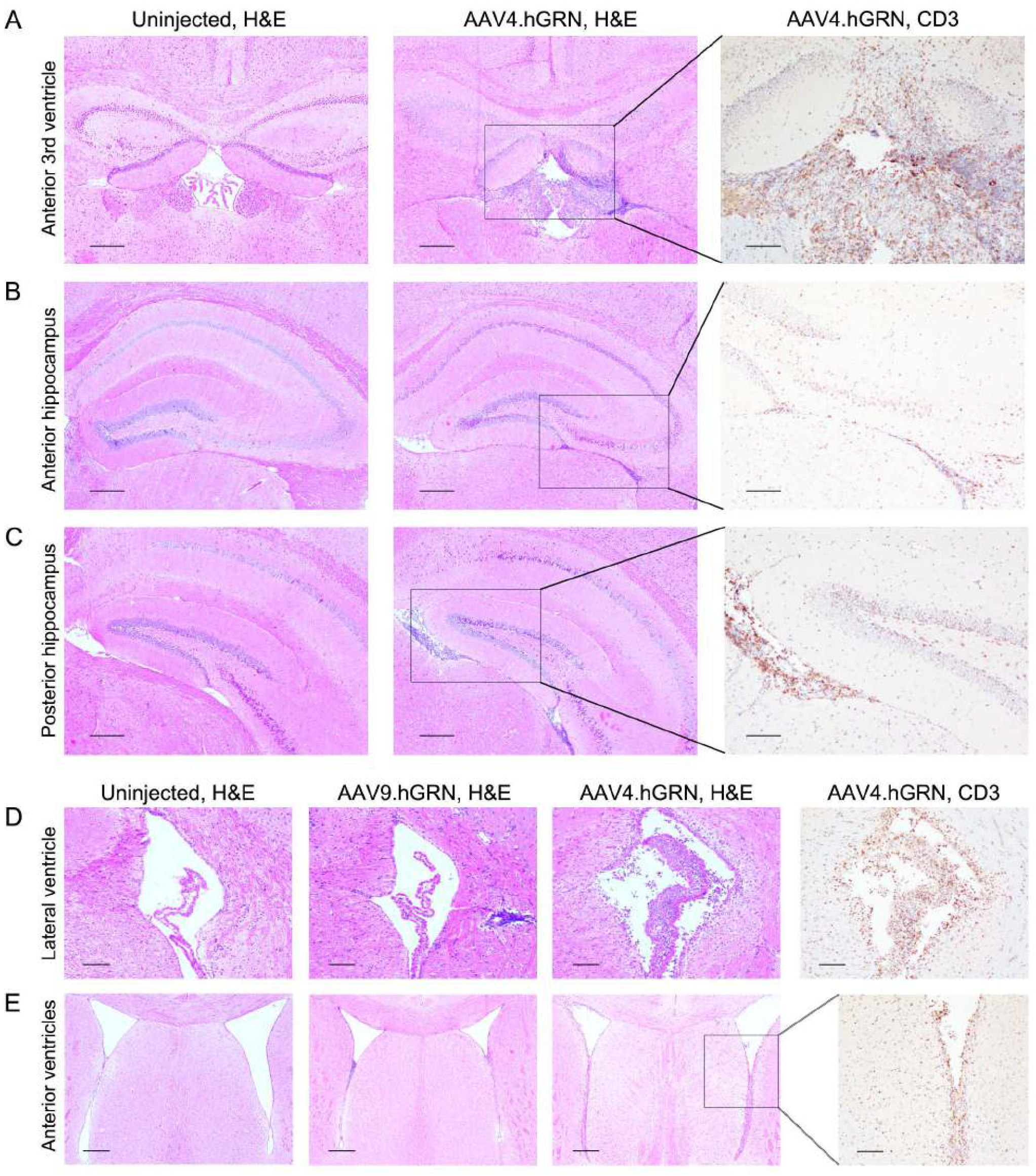
*Grn* null mice expressing hGRN delivered by an ependymal-targeting vector (AAV4) show an inflammatory response and ependymal hypertrophy. Mice were injected at 6.5-8 months of age in the right lateral ventricle with AAV4.*hGRN* and sacrificed 1 month post-injection. AAV4.*hGRN*-injected mice show a dense cellular infiltrate in the far anterior hippocampus (A, middle panel), anterior hippocampus (B, middle panel), and posterior hippocampus (C, middle panel) that is not present in uninjected age-matched controls (A-C, left panels) or in an AAV4.*eGFP*-injected littermate (data not shown). (Scale bars: 250 μm.) The infiltrate is highly positive for the T cell marker CD3 in all regions (A-C, right panels; scale bars: 100 μm). When compared to age-matched AAV9.*hGRN* injected mice, AAV4.*hGRN*-injected mice show unique choroidal and ependymal hyperplasia and thickening and dense T cell infiltration in the ipsilateral (D) and contralateral (data not shown) lateral ventricles (scale bars: 100 μm). AAV4.*hGRN*-injected mice also demonstrate ependymal hyperplasia and T cell infiltration in the far anterior ventricles adjacent to the striatum (E; scale bars: 250 μm) that are not apparent in uninjected or in AAV9.*hGRN*-injected mice. (*n* = 3 mice/group. 2 of the 3 AAV4.*hGRN*-injected mice were affected.)

We additionally observed marked ependymal and choroidal hypertrophy in the lateral ventricles adjacent to the hippocampus (**Fig. 8*D***) as well as the anterior 3rd ventricle (**Fig. 8*A***) and the ventricles adjacent to the striatum (**Fig. 8*E***), suggesting a direct effect on cellular proliferation by GRN. This hypertrophy was not observed with AAV9-mediated gene delivery (**Fig. 8*D*** and **Fig. 8*E***, second panels). As AAV9 does not efficiently transduce ependymal cells, these data indicate a hypertrophic effect on cells directly expressing the *hGRN* transgene. Cumulatively, our data suggest that *GRN* gene delivery may trigger harmful responses in the CNS, irrespective of the AAV serotype used for delivery.

## Discussion

In these studies we sought to test the safety and efficacy of AAV-mediated *GRN* expression in a *Grn* null mouse model, towards development of a gene replacement strategy for treatment of *GRN*-deficient FTD and NCL. To our surprise we found that AAV9-mediated *GRN* overexpression led to severe degeneration of the hippocampus in *Grn* null mice. The observed degeneration was markedly selective, with sparing of the cortex above, striatum anterior to, and thalamic structures inferior to the hippocampus, despite high GRN levels in these tissues. We also observed a cellular infiltrate primarily composed of T cells as well as perivascular cuffing preceding the onset of hippocampal degeneration and persisting until late stages of degeneration. In addition, we detected a T cell mediated immune response irrespective of the genetic background of the mouse injected, and irrespective of the AAV serotype used, as well as a direct hypertrophic effect of GRN. These data emphasize the need for caution in pursuing GRN delivery in the human CNS.

Neuronal loss in our study was region-selective, with marked hippocampal neurodegeneration, preceded by T cell infiltration. We considered various explanations for our findings. First, the choice of the AAV9 vector – known to transduce both neurons and glia and to achieve high levels of expression – might have resulted in toxic levels of transgene expression. However, eGFP delivered by AAV9 under the same conditions did not elicit a similar response. Additionally, *GRN* delivered by the ependymally-targeting AAV4 serotype also provoked a marked inflammatory response. These data suggest that the choice of serotype is not sufficient to explain our findings.

Second, the intraventricular delivery route chosen here may have triggered immunogenicity and downstream tissue destruction. In this respect, the recent reports by Arrant *et al*. demonstrating rescue in mouse models of FTD and NCL after intraparenchymal delivery of *Grn* are noteworthy (37, 38). Specifically, whether GRN was expressed in mice lacking one or both copies of *Grn*, this group did not report the dramatic hippocampal degeneration we found in our study. It is possible that the intraventricular route of transgene delivery exposes particular antigen-presenting cells to GRN, thus provoking the T cell infiltration and inflammatory response observed in our animals. Our intraventricular delivery strategy was based on considerations of eventual clinical translation for *GRN*-deficient human diseases, for which intraparenchymal delivery poses safety and feasibility issues that are less concerning in preclinical models. Moreover, intraventricular CNS delivery of many different transgenes has been successfully achieved in preclinical models, including in animals null for the therapeutic gene, arguing that the choice of delivery route is not the sole explanation for our findings. In mice, at doses and injection routes similar to ours, gene replacement strategies have consistently been safely achieved in null models; for instance, one group safely delivered the ATP-binding cassette transporter (ABCD1) gene in a mouse model of X-linked adrenoleukodystrophy (39). Thus, a final consideration is the direct effect of GRN itself.

While much of the literature in the >10 years since *GRN* mutations were first linked to neurodegeneration concerns the role of GRN as a neurotrophic factor (14, 16-18), *GRN* is also widely expressed in cell types ranging from epithelial cells to hematopoietic cells, macrophages, and T cells (12, 13). Early studies described its role in wound healing and regulation of inflammation, which while poorly understood involves an interplay between GRN, which is itself active, and its cleavage products, the granulins, which have opposing effects on a number of immune cell-mediated processes (13). Thus, the existing literature on GRN suggests that tight regulation of its expression, both on the transcriptional side and with respect to the protein cleavage events that generate daughter peptides, may be needed to avoid untoward immunological effects.

There is also extensive literature on involvement of GRN in cell growth and proliferation, both in normal development and in cancer (12, 13). Indeed, GRN is overexpressed and promotes cell growth in many tumors, including glioblastoma (51, 52). In this respect, our findings using AAV4-mediated delivery of *GRN* are noteworthy. In mice, AAV4 selectively targets ependymal cells and astrocytes in the subventricular zone (53). We therefore reasoned that AAV4-mediated delivery of *GRN* might be safer, as we would avoid directly targeting neurons and glia, while allowing for endogenous mechanisms of GRN uptake through its neuronal receptor sortilin-1. We found, however, that AAV4-mediated *GRN* delivery resulted in marked hypertrophy of the ependyma, suggesting direct effects by GRN on the targeted cells, again accompanied by T cell infiltration.

Our data should be compared to recent studies by Arrant *et al*., who reported reversal of phenotypes associated with GRN deficiency using AAV mediated *GRN* replacement therapy in *Grn* null mice, and no evidence of T cell infiltration or hippocampal degeneration. Despite differences in our approach (intraventricular versus intraparenchymal), vector (AAV9 or AAV4 versus AAV1), dose (higher in our study), and transgene (our vector does not have a Myc tag and is able to interact with sortilin), both we and Arrant *et al*. observe an immune response. In the case of Arrant *et al*., the response consists of profound microglial activation and MHCII presentation at the injection site, as well as antibodies to GRN detected in plasma. In our studies, by contrast, we observed T cell infiltration and destruction of neurons and astrocytes in the hippocampus. It is possible that, given a longer period of time, evidence of hippocampal degeneration would emerge in their studies. Indeed, Arrant *et al*. remarked that the upregulation of MHCII is typically associated with a T cell response, and speculated that a longer exposure could result in T cell mediated neuronal degeneration and functional deficits.

These findings raise concerns regarding how to safely translate these studies into humans. Specifically, our combined data suggest that GRN replacement could be highly immunogenic in NCL (*GRN* null) patients, while achieving the levels needed to reverse FTD will require use of vectors with high transduction efficiency. High levels of *GRN* expression delivered by transgene, in turn, may run risks of exposure to the tumorigenic growth effects of GRN suggested by our AAV4 data. In addition, these results suggest future pharmacological toxicity studies will be challenging, as over-expression of the human clinical product will be problematic in rodents.

In summary, while *GRN*-associated FTD and NCL may appear to be attractive targets for gene replacement therapy, our results suggest that concerns stemming from the identity and function of the transgene in question – *GRN* – are paramount in considerations of a path to human intervention. Specifically, work elucidating the molecular mechanisms by which GRN modulates inflammatory and growth responses, particularly in the CNS, are needed. In addition, our work highlights the potential for inflammatory and tumorigenic adverse effects with *GRN* overexpression, to which specific attention should be paid in preclinical models. More broadly, these findings call into question our current conception of the brain as an immune privileged organ, AAV as a safe vector for all CNS delivery, and maximization of gene expression as the goal of gene replacement therapy, as these are in tight interplay and should be carefully individualized to the specific therapy being developed.

## Materials and Methods

Detailed materials and methods can be found in SI Materials and Methods. Procedures involving animals were performed in accordance with Institutional Animal Care and Use Committee of the University of Pennsylvania. *Grn* null mice were generated as previously described (54) and provided to the University of Pennsylvania by the Nishihara laboratory at the University of Tokyo. AAV vectors were produced by the CHOP Research Vector Core. Specific tests used are described in the figure legends. Transfections were performed with Lipofectamine 2000 (Invitrogen). For cells and mouse tissues, protein was obtained using RIPA buffer with 0.2% PMSF and 0.1% protease inhibitors. GRN was quantified by human progranulin ELISA kit (Adipogen). For IHC, the following antibodies were used: anti-human GRN rabbit polyclonal antibody (developed by CNDR as previously described (15)), anti-NeuN rabbit polyclonal antibody (ABN78; Sigma-Aldrich), anti-GFAP rabbit polyclonal antibody (Z0334; Dako/Agilent), anti-Iba1 rabbit polyclonal antibody (Saf5299), anti-Ki67 rabbit monoclonal antibody (Ab16667; Abcam), anti-CD45R rat monoclonal antibody (RA3-6B2; Invitrogen,), and anti-CD3e rabbit monoclonal antibody (MA1-90582; Invitrogen). Autoantibody studies were performed as previously described (55).

## ACKNOWLEDGEMENTS

We thank Masugi Nishihara for providing us with *Grn* null animals and Virginia Lee for providing GRN antibody reagents. Sources of support for this project include the Pechenik Montague Award Fund (to ACP), the Benaroya Award Fund (to ACP), and the Brody Foundation (to DA).

## SUPPLEMENTARY MATERIALS

**Fig. S1.**
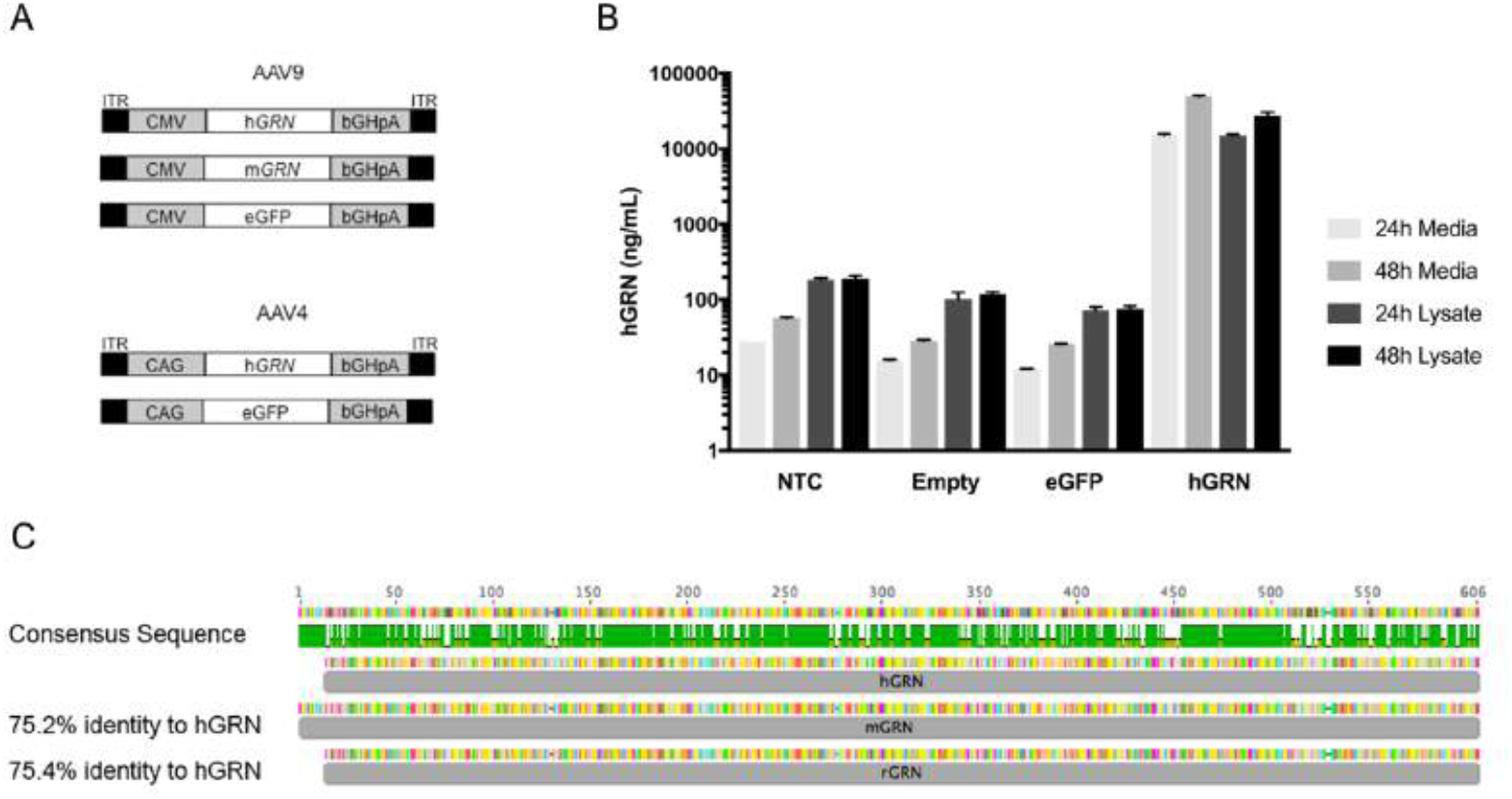
(A) Schematic of AAV transgene cassettes used in our experiments. For AAV9 vectors, the CMV promoter was used to drive human progranulin (*hGRN*), mouse progranulin (*mGrn*), or enhanced green fluorescent protein (*eGFP*), followed by the bovine growth hormone polyA (bGHpA), and flanked by AAV2 inverted terminal repeats (ITR). For AAV4 the CAG promoter was used to drive *hGRN* or *eGFP*, followed by the bovine growth hormone polyA (bGHpA), flanked by the AAV2 inverted terminal repeats (ITR). (B) Plasmid expression was validated by transfection of HEK293 cells (QBI) with lipofectamine 2000 and measuring hGRN or mGRN levels by ELISA in the media or lysate, 24 or 48 hours after transfection as indicated. Our expression plasmids are compared to non-transfected cells (NTC), cells transfected with the empty vector (5/TO) or *eGFP* transfected cells. (C) Schematic of hGRN, mGRN, and rat GRN (rGRN) protein consensus and alignments. rGRN and mGRN share 75.4% and 75.2% identity to hGRN respectively.

**Fig. S2.**
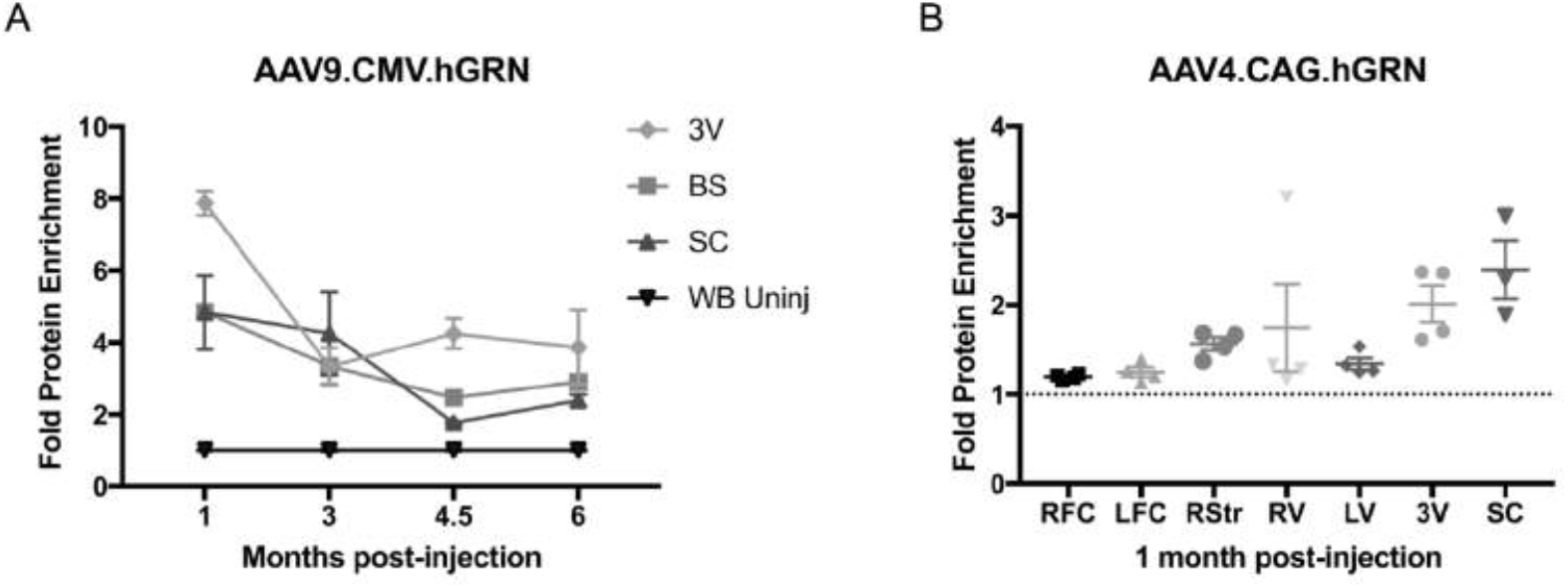
AAV9 and AAV4 mediate *hGRN* expression in *Grn* null mouse brain. *Grn* null mice were injected at 6-7.5 months of age with AAV9.*hGRN* (A) or AAV4.*hGRN* (B) in the right lateral ventricle, and brains were microdissected at various time points post-injection and hGRN levels measured by ELISA. (A) AAV9.*hGRN*-treated mice were sacrificed 1-, 3-, 4.5-, or 6 months post-injection. hGRN levels in the third peri-ventricular area (3V), brain stem (BS), spinal cord (SC) and uninjected homogenized whole brain (WB) are shown. Levels in these posterior regions are moderately sustained over time. (B) AAV4.*hGRN*-treated mice were sacrificed 1 month post-injection. hGRN levels in the right and left frontal cortex (RFC, LFC), right striatum (RStr), right peri-ventricular area (RV), 3V, and SC are shown, with uninjected homogenized WB represented by a dotted line. There is a small global increase in expression most pronounced in regions with a high ependymal content, such as RV, 3V and SC. *n* = 3 mice/group at each time point.

**Fig. S3.**
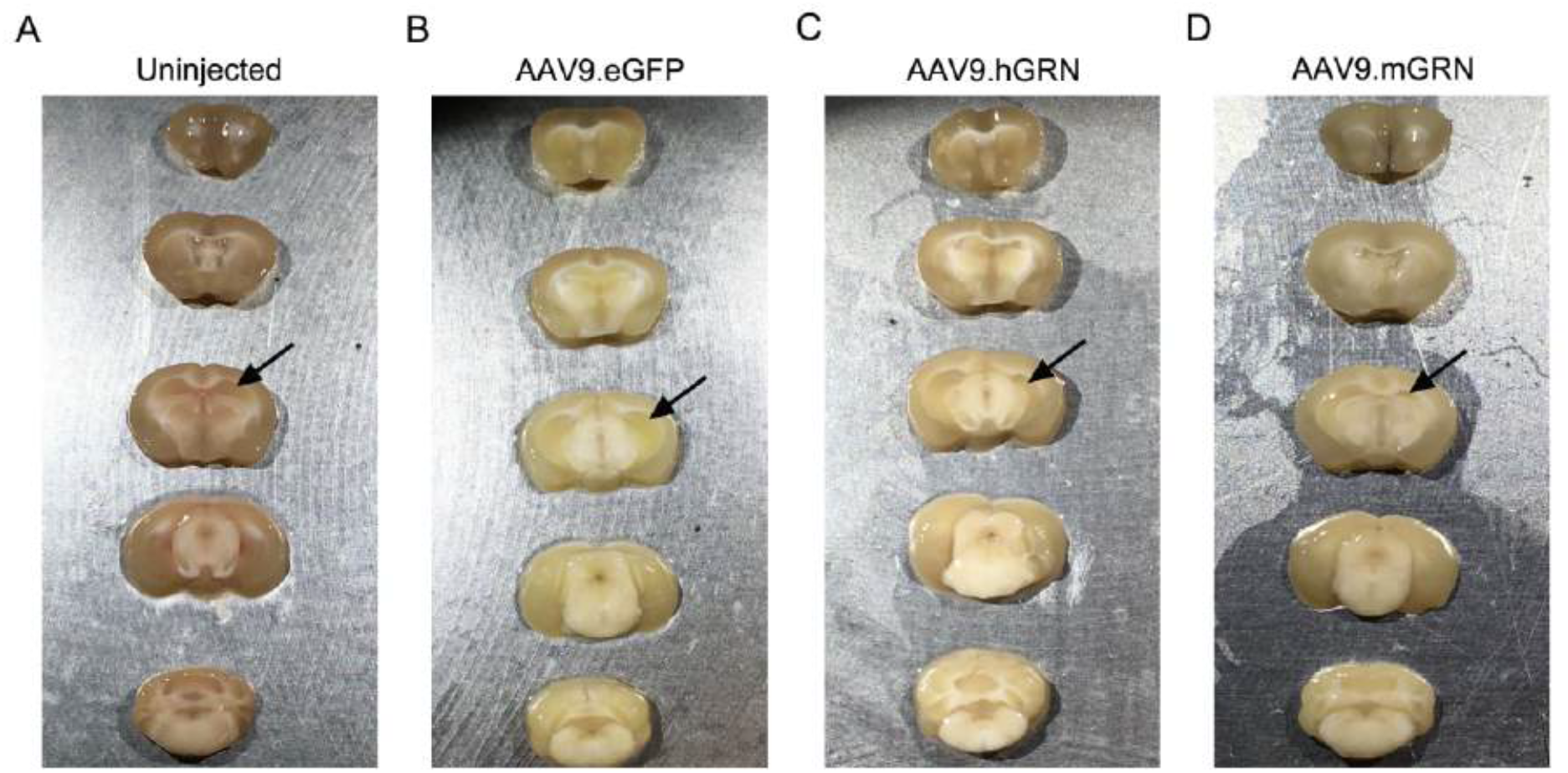
Gross morphological changes are observed in AAV9.*hGRN*- and AAV9.*mGrn*-injected *Grn* null mice. Mice were injected with AAV9.*hGRN*, AAV9.*mGrn*, or AAV9.*eGFP* at 6-7.5 months of age and sacrificed 6 months post-injection. Brains were cut into 2mm sections prior to microdissection. While uninjected (A) and AAV9.*eGFP*-injected (B) mice show no overt morphological differences, the injected (right) hemisphere of mice treated with AAV9.*hGRN* (C) and AAV9.*mGrn* (D) was noticeably smaller, specifically in the hippocampal region (arrows), with surrounding regions appearing unaffected. *n* =11 mice treated with AAV9.*hGRN (6 affected), n* = 3 mice treated with AAV9.*mGrn* (3 affected), *n* = 4 mice treated with AAV9.*eGFP* (0 affected), *n* = 4 mice uninjected (0 affected).

## EXPANDED MATERIALS AND METHODS

### Viral Vector Constructs

AAV serotype 9 and AAV serotype 4 were used for these studies. AAV9 vectors contained a cytomegalovirus (CMV) promoter, while AAV4 vectors contained a chicken ß actin (CBA) promoter. The human *GRN* cDNA (GenBank accession no. BC000324) was amplified from a Hek293 cell cDNA library and the mouse *Grn* cDNA (GenBank accession no. NM_008175) was cloned using gBlocks (IDT). The transgenes or an *eGFP* reporter gene were inserted into the G0347 pFB.AAV.CMV.bHGpA plasmid (Iowa Vector Core, Iowa City, IA) and were then used to produce AAV vectors by the Children’s Hospital of Philadelphia (CHOP) Research Vector Core (Philadelphia, PA). All constructs were verified by Sanger Sequencing (CHOP Napcore, Philadelphia, PA).

### Mice

Generation of progranulin-null (*Grn* null) mice on a C57BL/6J background through targeted disruption of the *Grn* gene has been reported previously (1). *Grn* null mice were provided to the University of Pennsylvania from the Nishihara laboratory at the University of Tokyo, and the colony was expanded and maintained. Male and female wildtype C57BL/6J (WT; Jackson labs, Bar Harbor, ME) and *Grn* null mice ages 6-12 months were used in these studies, as well as age-matched wildtype C57BL/6J controls. Mice were housed in a controlled temperature environment on a 12-hour light/dark cycle and were given free access to food and water. All animal studies were approved by the Institutional Animal Care and Use Committee of the University of Pennsylvania.

### Stereotaxic Delivery

Mice were deeply anaesthetized with isoflurane and immobilized in a stereotaxic frame (David Kopf instruments, Tujunga, CA) installed with both a digital stereotaxic control panel (Leica Biosystems, Buffalo Grove, IL) and microinjection robot (KD Scientific, Holliston, MA) for motorized injections. Mice were injected unilaterally in the posterior right lateral ventricle via a Hamilton syringe using the following coordinates: AP, −2.18 mm; ML, −2.9 mm; and DV, −3.5 mm (from bone) relative to bregma. Each mouse received 10 μL of vector at a concentration of 5e+12 vg/ml for a total dose of 5e+10 VG, infused at a rate of 0.5 μL/min with a 3 minute wait time post-infusion prior to withdrawal of the trochanter. Mice were injected between 6-7.5 months of age and sacrificed between 1 and 6 months post-injection.

### Mouse Brain Isolation

Mice were sacrificed at indicated ages by anesthetizing with a ketamine/xylazine/acepromazine mixture, followed by transcardial perfusion with 15mL of ice-cold 0.9% phosphate-buffered saline (PBS). Brains were quickly removed from the skull. Those used for fluorescent imaging studies were fixed whole in 4% paraformaldehyde overnight at 4°C followed by placement in a 30% sucrose/0.05% sodium azide solution for cryoprotection at 4°C. Those used for immunohistochemistry or for GRN quantification by ELISA were blocked into 2mm-thick coronal slices and then either fixed in 4% paraformaldehyde overnight at 4°C or microdissected and flash-frozen in liquid nitrogen, respectively.

### Protein Extraction

Frozen isolated brain tissues were weighed and transferred to a 500 μL Potter-Elvehjem dounce homogenizer (Sigma-Aldrich, Allentown, PA), and manually homogenized in 1% radioimmunoprecipitation buffer (RIPA; 50 mM Tris, 150 mM NaCl, 5mM EDTA, 0.5% sodium deoxycholate, 1% NP-40, 0.1% sodium dodecyl sulfate [SDS], pH 8.0) with 0.2% phenylmethane sulfonyl fluoride (PMSF) and 0.1% protease inhibitors (Penn Center for Neurodegenerative Research [CNDR], Philadelphia, PA). Homogenates were centrifuged at 21,380 RCF for 30 minutes and supernatant harvested, and protein concentration was measured by Pierce colorimetric BCA protein assay (ThermoFisher Scientific, Waltham, MA). GRN was quantified by human progranulin ELISA kit (Adipogen, San Diego, CA). 118 μg of total protein was plated per well, with samples run in duplicate. Plates were read on a Berthold LB941 TriStar vTI Multimode reader (Berthold Technologies, Bad Wildbad, Germany) at a wavelength of 450 nm and progranulin concentration calculated using the provided standard curve per kit instructions.

### Antibodies

Primary antibodies used for immunohistochemistry included the following: anti-human GRN (anti-hGRN) rabbit polyclonal antibody (0.01mg/mL; developed by CNDR, Philadelphia, PA as previously described (2)), anti-NeuN rabbit polyclonal antibody (0.5μg/mL; ABN78; Sigma-Aldrich, St. Louis, MO), anti-GFAP rabbit polyclonal antibody (0.58μg/mL; Z0334; Dako/Agilent, Santa Clara, CA), anti-Iba1 rabbit polyclonal antibody (0.25μg/mL; Saf5299; Wako, Richmond, VA), anti-Ki67 rabbit monoclonal antibody (dilution 1:1000; Ab16667; Abcam, Cambridge, UK), anti-CD45R rat monoclonal antibody (5μg/mL; RA3-6B2; Invitrogen, Carlsbad, CA), and anti-CD3e rabbit monoclonal antibody (dilution 1:150; MA1-90582; Invitrogen, Carlsbad, CA). Biotinylated goat anti-rabbit and goat anti-rat IgG secondary antibodies (Vector laboratories, Burlingame, CA) were used at a concentration of 1.5μg/mL in all cases except for anti-hGRN, for which secondary antibody concentration was 7.5μg/mL. In the autoantibody studies, biotinylated goat anti-mouse IgG secondary antibody (Vector laboratories, Burlingame, CA) was used at a concentration of 0.75μg/mL.

### Section preparation and immunohistochemistry

Fixed coronal brain slices were serially ethanol-dehydrated and paraffin-embedded. Blocks were sectioned coronally at 6 μm and incubated at 37-42°C overnight. Sections were deparaffinized in Xylene and rehydrated in serially dilute ethanol solutions, and were either stained with hematoxylin (ThermoScientific, Kalamazoo, MI) and eosin (Fisher Chemical, Waltham, MA) (H&E) or underwent immunohistochemical (IHC) staining, as follows: sections underwent deactivation of endogenous peroxidase with 5% hydrogen peroxide in methanol for 30 minutes as well as microwave antigen retrieval using antigen unmasking solution (Vector laboratories, Burlingame, CA), followed by washing in Tris buffer then blocking against nonspecific binding sites with 2% fetal bovine serum for 30 minutes at room temperature. Sections were then incubated in primary antibody overnight at 4°C in humidified chambers, blocked again, and incubated in biotinylated secondary antibody for one hour at room temperature. Sections were again blocked, followed by treatment for one hour with the VECTASTAIN ABC kit (Vector laboratories, Burlingame, CA) for avidin binding and peroxidation, then treated with Vector ImmPACT 3,3’-Diaminobenzidine (DAB) peroxidase substrate solution for detection (Vector laboratories, Burlingame, CA). Sections were counter-stained with hematoxylin and dehydrated prior to coverslipping. Brightfield images were taken on a Nikon 80i upright fluorescence microscope and analyzed with Nikon NIS-Elements AR Imaging software.

### Section preparation and fluorescence imaging

Fixed, cryoprotected brains were sectioned at 60 μm on a freezing microtome and stored at −20°C in a cryoprotectant solution (30% ethylene glycol, 15% sucrose, 0.05% sodium azide in phosphate buffered saline) until use. Sections were imaged using a DM6000B Leica microscope equipped with a Hamamatsu ORCA-Flash4.0 camera.

### Autoantibody detection

Studies were performed as previously described (3). Adult female Wistar rats were anesthetized and decapitated. Brains were removed and washed in 1x phosphate-buffered saline (PBS), then bisected sagitally and fixed in 4% paraformaldehyde in PBS at 4°C for one hour. Brains were then transferred to 40% sucrose in 0.1M PBS for 48 hours, followed by embedding in Tissue-Tek OCT compound embedding medium (Sakura Finitek, Torrance, CA), then snap-frozen in 2-methylbutane cooled with liquid nitrogen. Sections were cut sagitally at 7 μm, mounted on glass slides, and stored at −20°C until use. Endogenous peroxidase was quenched with 0.3% hydrogen peroxide in PBS for 15 minutes. Sections were blocked in 5% goat normal serum for one hour, after which serum samples were applied to sections at a dilution of 1:200 in 5% normal goat serum (Jackson ImmunoResearch Laboratories, West Grove, PA) and incubated overnight at 4°C in humidified chambers. Sections were incubated in biotinylated goat anti-mouse secondary antibody for 2 hours at room temperature, then treated for one hour with the VECTASTAIN Elite ABC kit (Vector laboratories, Burlingame, CA) for avidin binding and peroxidation. Slides were incubated for 30 seconds in 0.5% Triton X solution in PBS, then treated with Vector ImmPACT DAB peroxidase substrate kit (Vector laboratories, Burlingame, CA) for detection. Slides were counter-stained with 50% hematoxylin and dehydrated prior to coverslipping. Brightfield images were taken on a Nikon 80i upright fluorescence microscope and analyzed with Nikon NIS-Elements AR Imaging software.

## REFERENCES

1. Baker M, et al. (2006) Mutations in progranulin cause tau-negative frontotemporal dementia linked to chromosome 17. Nature 442(7105):916–919.

2. Cruts M, et al. (2006) Null mutations in progranulin cause ubiquitin-positive frontotemporal dementia linked to chromosome 17q21. Nature 442(7105):920–924.

3. Smith KR, et al. (2012) Strikingly different clinicopathological phenotypes determined by progranulin-mutation dosage. Am JHum Genet 90(6): 1102–1107.

4. Grossman M (2002) Frontotemporal dementia: a review. J Int Neuropsychol Soc 8(4):566–583.

5. Chen-Plotkin AS, et al. (2011) Genetic and clinical features of progranulin-associated frontotemporal lobar degeneration. Arch Neurol 68(4):488–497.

6. Canafoglia L, et al. (2014) Recurrent generalized seizures, visual loss, and palinopsia as phenotypic features of neuronal ceroid lipofuscinosis due to progranulin gene mutation. Epilepsia 55(6):e56–59.

7. Almeida MR, et al. (2016) Portuguese family with the co-occurrence of frontotemporal lobar degeneration and neuronal ceroid lipofuscinosis phenotypes due to progranulin gene mutation. Neurobiol Aging 41:200 e201–200 e205.

8. Kousi M, Lehesjoki AE, & Mole SE (2012) Update of the mutation spectrum and clinical correlations of over 360 mutations in eight genes that underlie the neuronal ceroid lipofuscinoses. Hum Mutat 33(1):42–63.

9. Capell A, et al. (2011) Rescue of progranulin deficiency associated with frontotemporal lobar degeneration by alkalizing reagents and inhibition of vacuolar ATPase. J Neurosci 31(5):1885–1894.

10. Cenik B, et al. (2011) Suberoylanilide hydroxamic acid (vorinostat) up-regulates progranulin transcription: rational therapeutic approach to frontotemporal dementia. J Biol Chem 286(18):16101–16108.

11. Lee WC, et al. (2014) Targeted manipulation of the sortilin-progranulin axis rescues progranulin haploinsufficiency. Hum Mol Genet 23(6):1467–1478.

12. Toh H, Chitramuthu BP, Bennett HP, & Bateman A (2011) Structure, function, and mechanism of progranulin; the brain and beyond. J Mol Neurosci 45(3):538–548.

13. Jian J, Konopka J, & Liu C (2013) Insights into the role of progranulin in immunity, infection, and inflammation. JLeukoc Biol 93(2):199–208.

14. Petkau TL, et al. (2010) Progranulin expression in the developing and adult murine brain. J Comp Neurol 518(19):3931–3947.

15. Chen-Plotkin AS, et al. (2010) Brain progranulin expression in GRN-associated frontotemporal lobar degeneration. Acta Neuropathol 119(1): 111–122.

16. Van Damme P, et al. (2008) Progranulin functions as a neurotrophic factor to regulate neurite outgrowth and enhance neuronal survival. J Cell Biol 181(1):37–41.

17. Gao X, et al. (2010) Progranulin promotes neurite outgrowth and neuronal differentiation by regulating GSK-3beta. Protein Cell 1(6):552–562.

18. Beel S, et al. (2017) Progranulin functions as a cathepsin D chaperone to stimulate axonal outgrowth in vivo. Hum Mol Genet 26(15):2850–2863.

19. Hu F, et al. (2010) Sortilin-mediated endocytosis determines levels of the frontotemporal dementia protein, progranulin. Neuron 68(4):654–667.

20. Minami SS, et al. (2014) Progranulin protects against amyloid beta deposition and toxicity in Alzheimer’s disease mouse models. Nat Med 20(10): 1157–1164.

21. Van Kampen JM & Kay DG (2017) Progranulin gene delivery reduces plaque burden and synaptic atrophy in a mouse model of Alzheimer’s disease. PLoS One 12(8):e0182896.

22. Van Kampen JM, Baranowski D, & Kay DG (2014) Progranulin gene delivery protects dopaminergic neurons in a mouse model of Parkinson’s disease. PLoS One 9(5):e97032.

23. Laird AS, et al. (2010) Progranulin is neurotrophic in vivo and protects against a mutant TDP-43 induced axonopathy. PLoS One 5(10):e13368.

24. Chitramuthu BP, Kay DG, Bateman A, & Bennett HP (2017) Neurotrophic effects of progranulin in vivo in reversing motor neuron defects caused by over or under expression of TDP-43 or FUS. PLoS One 12(3):e0174784.

25. Tauffenberger A, Chitramuthu BP, Bateman A, Bennett HP, & Parker JA (2013) Reduction of polyglutamine toxicity by TDP-43, FUS and progranulin in Huntington’s disease models. Hum Mol Genet 22(4):782–794.

26. Russell S, et al. (2017) Efficacy and safety of voretigene neparvovec (AAV2-hRPE65v2) in patients with RPE65-mediated inherited retinal dystrophy: a randomised, controlled, open-label, phase 3 trial. Lancet 390(10097):849–860.

27. Dunbar CE, et al. (2018) Gene therapy comes of age. Science 359(6372).

28. Mendell JR, et al. (2017) Single-Dose Gene-Replacement Therapy for Spinal Muscular Atrophy. N Engl J Med 377(18): 1713–1722.

29. Sands MS & Davidson BL (2006) Gene therapy for lysosomal storage diseases. Molecular therapy: the journal of the American Society of Gene Therapy 13(5):839–849.

30. Katz ML, et al. (2015) AAV gene transfer delays disease onset in a TPP1-deficient canine model of the late infantile form of Batten disease. Science translational medicine 7(313):313ra180.

31. Chen YH, Chang M, & Davidson BL (2009) Molecular signatures of disease brain endothelia provide new sites for CNS-directed enzyme therapy. Nat Med 15(10):1215–1218.

32. Monteys AM, Ebanks SA, Keiser MS, & Davidson BL (2017) CRISPR/Cas9 Editing of the Mutant Huntingtin Allele In Vitro and In Vivo. Molecular therapy: the journal of the American Society of Gene Therapy 25(1): 12–23.

33. Meyer K, et al. (2015) Improving single injection CSF delivery of AAV9-mediated gene therapy for SMA: a dose-response study in mice and nonhuman primates. Molecular therapy: the journal of the American Society of Gene Therapy 23(3):477–487.

34. Haurigot V, et al. (2013) Whole body correction of mucopolysaccharidosis IIIA by intracerebrospinal fluid gene therapy. J Clin Invest.

35. Mingozzi F & High KA (2017) Overcoming the Host Immune Response to Adeno-Associated Virus Gene Delivery Vectors: The Race Between Clearance, Tolerance, Neutralization, and Escape. Annual review of virology 4(1):511–534.

36. Vandamme C, Adjali O, & Mingozzi F (2017) Unraveling the Complex Story of Immune Responses to AAV Vectors Trial After Trial. Hum Gene Ther 28(11): 1061–1074.

37. Arrant AE, Onyilo VC, Unger DE, & Roberson ED (2018) Progranulin Gene Therapy Improves Lysosomal Dysfunction and Microglial Pathology Associated with Frontotemporal Dementia and Neuronal Ceroid Lipofuscinosis. J Neurosci 38(9):2341–2358.

38. Arrant AE, Filiano AJ, Unger DE, Young AH, & Roberson ED (2017) Restoring neuronal progranulin reverses deficits in a mouse model of frontotemporal dementia. Brain 140(5):1447–1465.

39. Gong Y, et al. (2015) Adenoassociated virus serotype 9-mediated gene therapy for x-linked adrenoleukodystrophy. Molecular therapy: the journal of the American Society of Gene Therapy 23(5):824–834.

40. Liu G, Martins I, Wemmie JA, Chiorini JA, & Davidson BL (2005) Functional correction of CNS phenotypes in a lysosomal storage disease model using adeno-associated virus type 4 vectors. J Neurosci 25(41):9321–9327.

41. Dodge JC, et al. (2010) AAV4-mediated expression of IGF-1 and VEGF within cellular components of the ventricular system improves survival outcome in familial ALS mice. Molecular therapy: the journal of the American Society of Gene Therapy 18(12):2075–2084.

42. Ahmed Z, et al. (2010) Accelerated lipofuscinosis and ubiquitination in granulin knockout mice suggest a role for progranulin in successful aging. Am J Pathol 177(1):311–324.

43. Wils H, et al. (2012) Cellular ageing, increased mortality and FTLD-TDP-associated neuropathology in progranulin knockout mice. J Pathol 228(1):67–76.

44. Ghoshal N, Dearborn JT, Wozniak DF, & Cairns NJ (2012) Core features of frontotemporal dementia recapitulated in progranulin knockout mice. Neurobiol Dis 45(1):395–408.

45. Sofroniew MV & Vinters HV (2010) Astrocytes: biology and pathology. Acta Neuropathol 119(1):7–35.

46. Lancaster E, et al. (2011) Antibodies to metabotropic glutamate receptor 5 in the Ophelia syndrome. Neurology 77(18): 1698–1701.

47. Lancaster E & Dalmau J (2012) Neuronal autoantigens--pathogenesis, associated disorders and antibody testing. Nat Rev Neurol 8(7):380–390.

48. Foust KD, et al. (2009) Intravascular AAV9 preferentially targets neonatal neurons and adult astrocytes. Nat Biotechnol 27(1):59–65.

49. Davidson BL, et al. (2000) Recombinant adeno-associated virus type 2, 4, and 5 vectors: transduction of variant cell types and regions in the mammalian central nervous system. Proceedings of the National Academy of Sciences of the United States of America 97(7):3428–3432.

50. Hudry E, et al. (2013) Gene transfer of human Apoe isoforms results in differential modulation of amyloid deposition and neurotoxicity in mouse brain. Science translational medicine 5(212):212ra161.

51. Liau LM, et al. (2000) Identification of a human glioma-associated growth factor gene, granulin, using differential immuno-absorption. Cancer Res 60(5):1353–1360.

52. Menges CW, et al. (2010) A Phosphotyrosine Proteomic Screen Identifies Multiple Tyrosine Kinase Signaling Pathways Aberrantly Activated in Malignant Mesothelioma. Genes Cancer 1(5):493–505.

53. Liu G, Martins IH, Chiorini JA, & Davidson BL (2005) Adeno-associated virus type 4 (AAV4) targets ependyma and astrocytes in the subventricular zone and RMS. Gene therapy 12(20):1503–1508.

54. Kayasuga Y, et al. (2007) Alteration of behavioural phenotype in mice by targeted disruption of the progranulin gene. Behav Brain Res 185(2): 110–118.

55. McCracken L, et al. (2017) Improving the antibody-based evaluation of autoimmune encephalitis. Neurol Neuroimmunol Neuroinflamm 4(6):e404.

## References

1. Kayasuga Y, et al. (2007) Alteration of behavioural phenotype in mice by targeted disruption of the progranulin gene. Behav Brain Res 185(2): 110–118.

2. Chen-Plotkin AS, et al. (2010) Brain progranulin expression in GRN-associated frontotemporal lobar degeneration. Acta Neuropathol 119(1): 111–122.

3. McCracken L, et al. (2017) Improving the antibody-based evaluation of autoimmune encephalitis. Neurol Neuroimmunol Neuroinflamm 4(6):e404.

